# The promoter of *Zymomonas mobilis* respiratory NADH dehydrogenase (*ndh*) is induced by oxygen

**DOI:** 10.1101/2025.01.06.631551

**Authors:** Marta Rubina, Inese Strazdina, Reinis Rutkis, Uldis Kalnenieks

## Abstract

Expression of the genes of engineered green fluorescent protein and the *Zymomonas mobilis* native malic enzyme from plasmid vectors under the *Z. mobilis* respiratory NADH dehydrogenase promoter (Pndh) was strongly enhanced by aeration, both in the wild type Zm6 and its respiratory-deficient mutant derivative Zm6-*ndh* backgrounds. Pndh in aerobically growing cultures was activated by about an order of magnitude relative to non-aerated control. Its induction approached the maximum level already at moderate aeration (1-5% oxygen saturation in the medium). The strength of Pndh under aerobic conditions was comparable to, or even surpassed that of the strong *Z. mobilis* native promoter of glyceraldehyde-3-phosphate dehydrogenase. Although the mechanism of its oxygen-dependent induction is not known, Pndh might serve as a versatile inducible promoter for *Z. mobilis* metabolic engineering.

**Highlights:**

- Aeration activates the *Z. mobilis ndh* promoter Pndh by about an order of magnitude
- Under aerobic conditions Pndh ranks among the strongest native *Z. mobilis* promoters
- Pndh is induced already at low oxygen saturation range (1-5%)

---

The ethanologenic alpha-proteobacterium *Zymomonas mobilis* bears several outstanding traits of biotechnological interest: an extremely high catabolic rate, tolerance to high ethanol and sugar concentrations, broad pH range of production, GRAS status (for a recent review, see Braga et al., 2021). Yet, in order to turn it into a versatile industrial platform microorganism, more tools are needed to modify its central metabolism and to introduce novel heterologous pathways (Behrendt et al., 2024; Kalnenieks et al., 2020). Promoters, in particular inducible promoters, are among the key parts of any metabolic engineering toolkit, as they ensure a flexible transcription-level control, and hence, contribute to the regulation of the activity of biosynthetic pathways. Although recently a systematic investigation of *Z. mobilis* promoter properties has been reported (Yang et al., 2019), the selection of well-characterized inducible promoters for application in this bacterium is still small. A few studies have used IPTG and tetracycline-inducible systems (Liu et al., 2020; Hu et al., 2023; Frohwitter et al., 2024). However, native inducible promoters derived from *Z. mobilis* are not in use up to date.

Here we present evidence that the *Z. mobilis* Type II respiratory NADH dehydrogenase (*ndh*) promoter (Pndh) is activated by aeration. Compared with other types of promoter induction, oxygen-dependent induction is advantageous for bioprocess control (Baez et al., 2014), because it does not require addition of chemical inducers that are (i) relatively expensive for large-scale use, and (ii) may interfere with the producer strain metabolism. Oxygenation of the culture can be easily controlled, and furthermore, its variation does not affect the composition of the growth medium, thus simplifying the product recovery.

To explore the oxygen effect on the Pndh activity, we transformed the wild type strain Zm6 (ATCC 29191) and its mutant derivative, the NADH dehydrogenase knock-out strain Zm6-*ndh* (Kalnenieks et al., 2008) with a reporter system, the plasmid vector pBBR:Pndh-eGFP. The rationale behind exploring the Zm6-*ndh* genetic background was its low oxygen consumption rate that would allow to use an oxygen-inducible promoter without compromising the intracellular NADH/NAD balance. That would be of particular importance, should a reduced compound (e.g. ethanol) be the target product. The strains, plasmid vectors, primers, used in the study, as well as the amplified promoter regions are presented in the Supplement files I – III. All plasmid and strain construction procedures were same as in Strazdina et al. (2023).

It is known that cells may experience oxidative stress when eGFP is expressed (Ganini et al., 2017) that might affect also the promoters under study. Furthermore, the eGFP fluorescence itself depends on oxygen, which somewhat complicates its use as a reporter for oxygen-dependent promoter induction (Vordermark et al., 2001). Therefore, we validated the results obtained with eGFP by using an alternative indicator, the *Z. mobilis* native malic enzyme (the decarboxylating NAD-dependent malate dehydrogenase; Mae). The strong constitutive promoter of *Z. mobilis* glyceraldehyde-3-phosphate dehydrogenase (Pgap) (Yang et al., 2019; Behrendt et al., 2022) was taken as a reference to characterize the relative strength of Pndh, with both eGFP and Mae.

Batch cultivations at 30 °C were performed on complex medium with 50 g/L glucose, supplemented with yeast extract and mineral salts (Kalnenieks et al., 1993). Cultures were grown either in closed 30 mL Falcon tubes containing 30 mL culture volume in a thermostat (microaerobic conditions), or in 500 mL shaken flasks with 60 mL culture volume on a shaker at 170 r.p.m. (aerobic conditions). Batch fermentations with pO_2_ control were carried out in a 0.5 L tabletop fermentor Sartorius Stedim Biotech, model Biostat Q Plus, using 3 working volumes in parallel. Cells were harvested 8 hours after inoculation, when the cultures were entering their early stationary growth phase. Bacteria were sedimented, washed and disintegrated by ultrasonication as in (Strazdina et al., 2012). After sedimentation of unbroken cells the cell-free extract was either (i) subject to eGFP fluorescence measurement with a preceding 4 h aeration on a shaker at 170 r.p.m. (for the strains, carrying plasmid vectors with *egfp*), in order to develop eGFP fluorescence in the cells grown under microaerobic conditions (Vordermark et al., 2001), or (ii) its membrane fraction was sedimented by ultracentrifugation (Kalnenieks et al., 1993) and the supernatant used for monitoring of malic enzyme activity (for the strains carrying plasmid vectors with *mae* and for the parent strains). The eGFP fluorescence was measured with a FluoroMax-3 spectrofluorimeter (Jobin–Yvon) at 475 nm excitation wavelength and 510 nm emission wavelength. The Mae activity was measured in the direction of malate oxidative decarboxylation, monitoring NAD reduction at 340 nm, as in Bringer-Meyer and Sahm (1989). Data represent means of three biological repeats, with error bars showing standard deviation.

The eGFP fluorescence revealed strong activation of Pndh under aerobic growth conditions (Fig. 1). In the strain Zm6, the Pndh was induced by a factor of more than 50, while in the Zm6-*ndh* induction factor was close to 20. The constitutive reference promoter Pgap, however, also showed an aerobic induction by a factor of 3, that was not seen with Mae (Fig. 2). That might partly be due to an incomplete development of the eGFP fluorescence in the cell-free extracts of the microaerobic cultures, despite their aerobic incubation. Nevertheless, even if that was the case, these eGFP data clearly indicated aerobic induction of Pndh – the aerobic increase of fluorescence with Pndh exceeded that with Pgap by about an order of magnitude in both strain backgrounds.

**Fig. 1.**
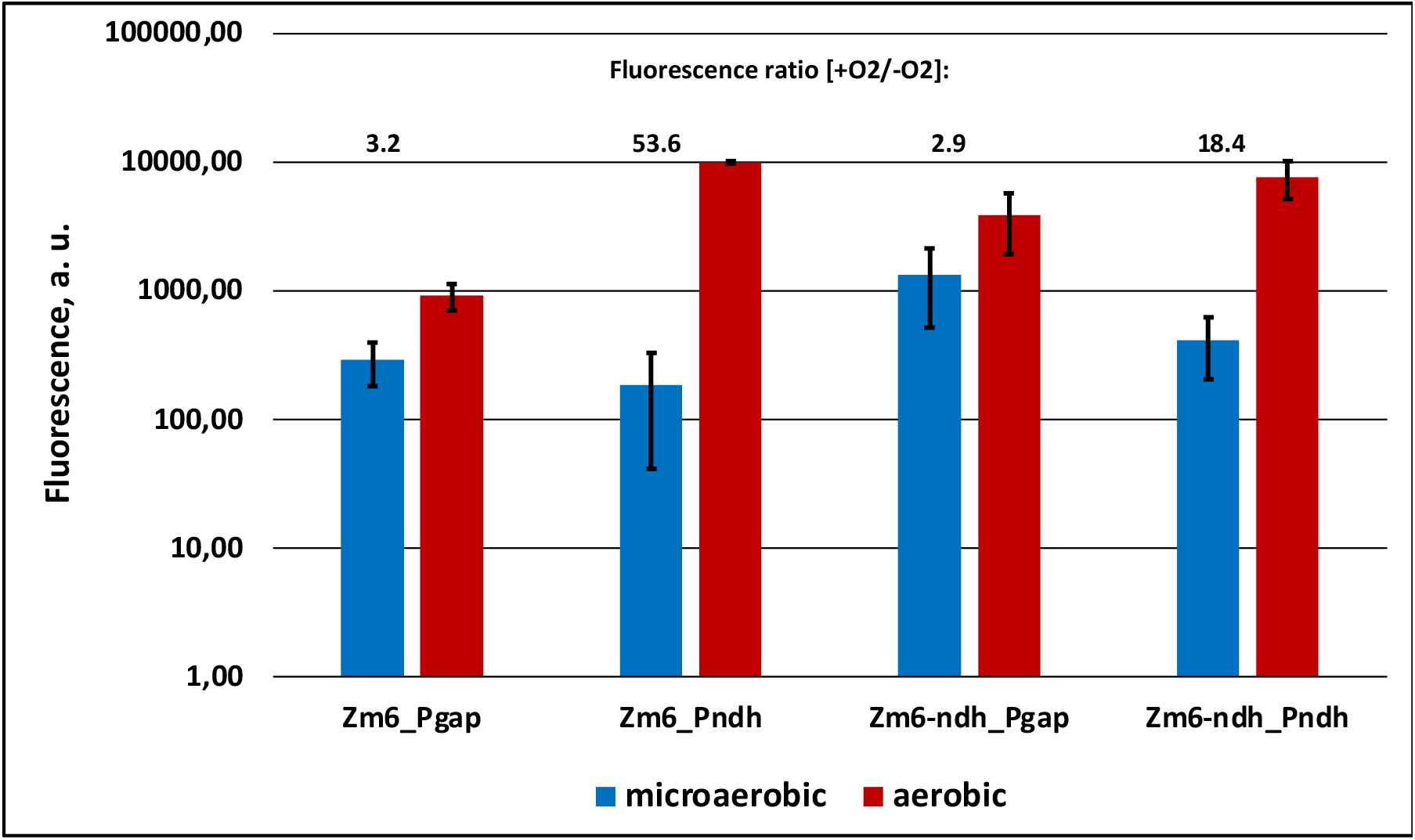
Effect of culture aeration on the expression of eGFP from the pBBR plasmid vector under gap promoter (Pgap) and under ndh promoter (Pndh) in the wild-type background, Zm6, and in the *ndh*-negative background, Zm6-*ndh*). First, for the cell extract of each strain, the recorded fluorescence value was normalized per milligram of protein. The normalized fluorescence of the strain Zm6_pBBR-eGFP was taken as the baseline, implying that all of the fluorescence exceeding that level in the other eGFP-bearing strains resulted from the activity of either Pgap or Pndh. Accordingly, the fluorescence values presented here show the difference between the normalized fluorescence of any of the promoter-carrying strains and the normalized fluorescence of the strain Zm6_pBBR-eGFP.

**Fig. 2.**
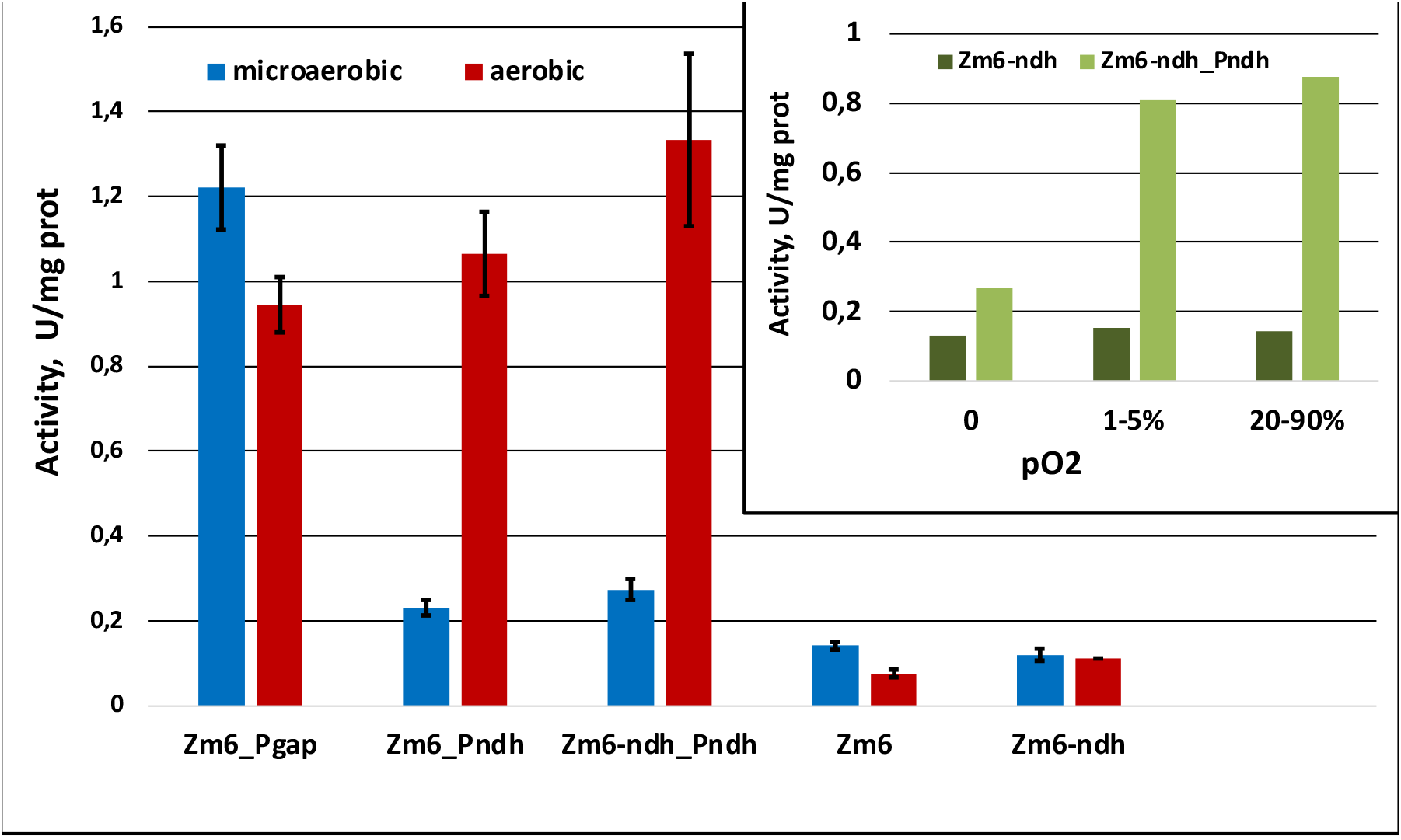
Effect of culture aeration on the overexpression of *Z. mobilis* Mae activity from the pBBR plasmid vector under gap promoter (Zm6_Pgap), under ndh promoter (against the wild-type background, Zm6_Pndh, and against the *ndh*-negative background, Zm6-*ndh*_Pndh), and solely from the chromosome under the *mae* native promoter in both strains. **Inset:** the effect of oxygen partial pressure in the culture on the Mae expression during controlled batch cultivations in fermentor. Fermentations with Zm6-*ndh* or Zm6-*ndh*_Pndh were carried out in 3 parallel working volumes, each with 200 mL of culture, with gassing at 100 mL min^-1^ and mixing at 200 r.p.m. Zero pO_2_ was established by gassing with pure nitrogen gas; the interval 20-90% corresponded to pO_2_ change during the 8 h of growth, when culture was gassed with air; pO_2_ was kept within 1-5% by manual regulation of the air / N_2_ proportion in the fermentor influx gas in the course of culture growth.

Also, with Mae aerobic induction of Pndh was observed (Fig. 2). The Mae activity ratio, calculated as: ([the vector-carrying strain – wild type]_aerobically_ / [the vector-carrying strain – wild type]_anaerobically_) showed induction of Pndh by a factor of 11 in Zm6 and by a factor of 6 (cultivation in the fermentor, the Fig.2 Inset) to 8 (cultivation in shaken flasks) in the Zm6-*ndh* background. Although this was a slightly lower level of induction, than seen with eGFP (in particular, for the Zm6 background), both reporter systems clearly indicated (i) aerobic induction of Pndh taking place, and (i) a high strength of Pndh under aerobic conditions, comparable to, or even surpassing the strength of Pgap.

It is well established that several *E. coli* oxygen-responsive promoters are being induced by transition from aerobic to anaerobic environment (Unden and Bongaerts 1997; Wichmann et al., 2023). Not so many are those induced by the opposite transition, like the Pndh in the present study. To the best of our knowledge, the *E. coli* SoxRS regulon is the only bacterial oxygen-inducible system with a potential biotechnological relevance that has been reported in the literature (for a review, see DeBaets et al., 2024). Baez et al. (2014) used *soxS* promoter for oxygen-dependent overexpression of the GFP. Its induction with oxygen raised the level of GFP expression in *E. coli* by about an order of magnitude, which is comparable to the effect observed in the present study. However, full induction of *soxS* promoter required as high as 300% saturation of the growth medium with oxygen. In contrast, the *Z. mobilis* Pndh was stimulated already at low pO_2_. A near-maximum value of its induction was reached in the range of 1 – 5% of oxygen saturation (Fig. 2, inset). Obviously, sensitivity of an oxygen-responsive promoter to low oxygen concentrations is an advantage. Low pO_2_ in the culture medium can be easier established and maintained, and creates no harmful physiological side effects. While, according to Baez et al. (2014), *E. coli* can survive 300% oxygen saturation levels quite well, growth of *Z. mobilis* is inhibited under vigorous aeration (Bringer et al., 1984), in part due to the redox imbalance of its catabolism leading to accumulation of the inhibitory metabolite acetaldehyde. pO_2_ in the range of 1 – 5%, however, does not significantly affect the growth of the *Z. mobilis* wild type. Furthermore, the growth of the strain Zm6-*ndh* is not inhibited also at higher pO_2_ (Kalnenieks et al., 2008). We conclude, therefore, that Pndh is a promising regulable promoter for *Z. mobilis* metabolic engineering applications.

Interestingly, the NADH:CoQ_1_ oxidoreductase activity in *Z. mobilis* membrane is less sensitive to oxygenation, than might be anticipted from our Pndh reporter system data. It tends to be slightly upregulated (approx. 1.5-3 times; unpublished observations) in aerobically growing *Z. mobilis* cells during the exponential phase. At later growth stage the aerobic *ndh* expression is reported to fall even below its anaerobic level (Yang et al., 2009). Also, the oxygen uptake rate of whole cells, and the membrane NADH oxidase activity generally is lower in aerobic cultures (Bringer et al., 1984; Strazdina et al., 2012). Interpretation of these observations is complicated by the fact that almost nothing is known about the redox-dependent regulatory systems in *Z. mobilis*. Its genome does not contain homologues to the sequences encoding, e.g., Fnr, ArcAB, or SoxRS regulators of *E. coli*. Understanding the regulation that underlies the *Z. mobilis* Pndh induction by oxygen, and, at the same time, the low oxygen-sensitivity of its respiratory chain will need further research, and might reveal some novel bacterial redox sensory mechanisms.

## Supporting information

Supplemental files I - III

## Acknowledgements

This research was funded by the projects Y5-AZ20-ZF-N-270 and Y9-B040-ZF-N-270 from the University of Latvia

