## Supplemental files I - III for "The promoter of *Zymomonas mobilis* respiratory NADH dehydrogenase (*ndh*) is induced by oxygen"

**Supplement I**

**Plasmids and strains used in the study**

**Plasmid/Strain	Characteristics	Source**

pBBR-eGFP	pBBR1MCS-2 derivative with an 0.72 kb insert of the	Strazdina et al., 2023

eGFP coding sequence in the MCS between EcoRI and

HindIII restriction sites

pBBR:Pndh-eGFP	pBBR-eGFP derivative with an 0.26 kb insert, containing	Strazdina et al., 2023

the promoter region of the Type II respiratory NADH

dehydrogenase (*ndh*, gene ZZ6_0213), upstream of the

eGFP coding sequence in the MCS between BamHI and

EcoRI restriction sites

pBBR:Pndh-Mae	pBBR:Pndh-eGFP derivative with a 1.69 kb insert of synthetic	present work

*Z. mobilis* malic enzyme (*mae*; gene ZZ6_1220) coding

sequence replacing the eGFP coding sequence downstream

the *ndh* promoter region, in the MCS between EcoRI and

HindIII restriction sites

pBBR:Pgap-eGFP	pBBR-eGFP derivative with an 0.3 kb insert, containing	present work

the promoter region of the Type I glyceraldehyde-3-

phosphate dehydrogenase (*gap*, gene ZZ6_1034), upstream

of the eGFP coding sequence in the MCS between BamHI

and EcoRI restriction sites

pBBR:Pgap-Mae	pBBR:Pgap-eGFP derivative with a 1.69 kb insert of synthetic	present work

*Z. mobilis* malic enzyme (*mae*; gene ZZ6_1220) coding

sequence replacing the eGFP coding sequence downstream

the *gap* promoter region, in the MCS between EcoRI and

HindIII restriction sites

Zm6	wild type, parent strain	ATCC 29191

Zm6-*ndh*	Zm6 with a Cm^r^ (chloramphenicol acetyltransferase gene)	Kalnenieks et al., 2008

inserted in the AgeI site in the chromosomal ORF of its *ndh*

Zm6_ pBBR-eGFP	Zm6 transformed with plasmid pBBR-eGFP	Strazdina et al., 2023

Zm6_ Pndh-eGFP	Zm6 transformed with plasmid pBBR:Pndh-eGFP	Strazdina et al., 2023

Zm6-*ndh*_ Pndh-eGFP	Zm6-*ndh* transformed with plasmid pBBR:Pndh-eGFP	present work

Zm6_Pndh-Mae	Zm6 transformed with plasmid pBBR:Pndh-Mae	present work

Zm6-*ndh*_ Pndh-Mae	Zm6-*ndh* transformed with plasmid pBBR:Pndh-Mae	present work

Zm6_ Pgap-eGFP	Zm6 transformed with plasmid pBBR:Pgap-eGFP	present work

Zm6-*ndh*_ Pgap-eGFP	Zm6-*ndh* transformed with plasmid pBBR:Pgap-eGFP	present work

Zm6_Pgap-Mae	Zm6 transformed with plasmid pBBR:Pgap-Mae	present work

Zm6-*ndh*_ Pgap-Mae	Zm6-*ndh* transformed with plasmid pBBR:Pgap-Mae	present work

**Supplement II**

**Primers used in the study**

**Primer name	Sequence with engineered restriction site underlined**

Pndh_BamHI_fwr	GGTTTGTGGATCCATAACCCATATTGACGG

Pndh_EcoRI_rev	GACATGAATTCCCTCTATTCTCTTCCAAGC

Pgap_BamHI_fwr	TTTTGGATCCGTTCGATCAACAACCCGAATC

Pgap_EcoRI_rev	CCCCCCGAATTCGTTTATTCTCCTAACTTATT

**Supplement III**

**The sequences of the amplified promoter regions**

***ndh*:**

**ATAACCCATATTGACGGGAAAAGAAGCCAAAGGTCAATCAGAGAAAGCGCGCTTTAAACAAAGCCCATGAGGATGATCAAAGACAGAGACCGGATAAGGGCAAAGATAGTTTCACAGTAAGCAATAAAAACTTTCACTCTGACTTATAATTTATGTGAAACAGATTTGAATGTTCTTCGTTAGTTACAGCAAAAGGTAATTGAGGCGGCTCGCAAACCACCACGCCAAAGGGGGCTGGGTCGCTTGGAAGAGAATAGAGGTTTCAA**

***gap*:**

**GTTCGATCAACAACCCGAATCCTATCGTAATGATGTTTTGCCCGATCAGCCTCAATCGACAATTTTACGCGTTTCGATCGAAGCAGGGACGACAATTGGCTGGGAACGGTATACTGGAATAAATGGTCTTCGTTATGGTATTGATGTTTTTGGTGCATCGGCCCCGGCGAATGATCTATATGCTCATTTCGGCTTGACCGCAGTCGGCATCACGAACAAGGTGTTGGCCGCGATCGCCGGTAAGTCGGCACGTTAAAAAATAGCTATGGAATATAGTAGCTACTTAATAAGTTAGGAGAATAAAC**
